# Human-calibrated automated intelligibility evaluation of speech neuroprostheses

**DOI:** 10.64898/2026.10.27.762196

**Authors:** Aparna Srinivasan, Maitreyee Wairagkar, Ramie Wakeb, Dana Farias, Lee M. Miller, David M. Brandman, Sergey D. Stavisky

**Affiliations:** Department of Neurological Surgery, University of California Davis, Davis, CA, USA; Biomedical Engineering Graduate Group, University of California Davis, Davis, CA, USA; Department of Neurobiology, Physiology and Behavior, University of California Davis, Davis, CA, USA; Department of Physical Medicine and Rehabilitation, University of California Davis Medical Center, Sacramento, CA, USA; Center for Mind and Brain, University of California Davis, Davis, CA, USA; Department of Otolaryngology - Head and Neck Surgery, University of California Davis, Davis, CA, USA

**Author notes:** co-senior authors (Lee M. Miller, David M. Brandman, Sergey D. Stavisky).

**Keywords:** Brain-computer interfaces, speech neuroprostheses, automatic speech recognition, low-rank adaptation

## Abstract

**Objective:** Brain-to-voice neuroprostheses can synthesize speech from the neural activity of people who have lost the ability to speak, enabling more natural and expressive communication. Assessing the intelligibility of synthesized speech, however, remains challenging because there is no ground-truth acoustic signal. The field instead relies on human transcription, which is costly and difficult to scale or compare across studies. Automatic speech recognition (ASR) offers a potential alternative, but state-of-the-art ASR models are optimized for transcription accuracy on typical fluent speech and may not reflect how humans perceive atypical synthesized speech.

**Approach:** We collected human transcriptions for ∼14.5 hours of synthesized speech from intracortical brain-to-voice decoders for one participant (yielding ∼72 hours of transcribed speech). We used the human-transcribed synthesized speech to fine-tune Whisper-large-v3 with low-rank adaptation (LoRA) so that its word error rate (WER) approximates human WER. We then evaluated the model on held-out sentences.

**Main results:** Fine-tuning increased the Pearson correlation between model WER and human WER from 0.62 to 0.76. The fine-tuned model’s WER closely tracked human WER across evaluation sets, including speech synthesized by unseen brain-to-voice decoders. Close agreement with human WER was achieved with only 1.5 hours of synthesized speech.

**Significance:** These results demonstrate that human-calibrated ASR can provide a scalable measure of brain-to-voice intelligibility that reflects human transcription more closely than zero-shot ASR does.

## Introduction

Speech brain-computer interfaces (BCIs) have the potential to restore communication for people who have lost the ability to speak due to neurological disease or injury [1,2]. Intracortical BCIs have demonstrated increasingly accurate decoding of attempted speech into text, enabling rapid communication by people with severe paralysis and dysarthria [3–5]. Recently, brain-to-voice BCIs have demonstrated real-time synthesis of attempted speech directly from neural activity, including the ability to convey aspects of natural speech such as intonation and emphasis [6,7].

The emergence of brain-to-voice BCIs introduces a distinct challenge for evaluating speech intelligibility. Brain-to-text BCIs can be evaluated against a known ground-truth text using a standard measure of transcription accuracy called word error rate (WER) [3,4,8–12]. In contrast, brain-to-voice BCIs generally lack ground-truth acoustic recordings corresponding to the synthesized speech. This makes human transcription a necessary component of intelligibility evaluation: listeners transcribe the synthesized speech, and their WER provides a behavioral measure of how intelligible the output is [6–8]. Human transcription, however, is time-consuming, costly, difficult to scale, and subject to inter-listener variability. Further, the transcriptions cannot be produced rapidly or frequently enough to serve as a loss function in training brain-to-voice decoders.

Automatic speech recognition (ASR) provides an appealing alternative to human transcription and has increasingly been used to automate intelligibility evaluation of both synthesized and atypical speech. In text-to-speech research, ASR-derived metrics have been used as an objective and scalable measure of intelligibility, reducing reliance on human listening tests [13–15]. ASR models have also been used to evaluate dysarthric speech, either by comparing ASR transcriptions with the intended text [16] or as feature extractors to predict listener effort ratings [17]. For speech synthesized by BCIs, our previous work used zero-shot ASR to transcribe brain-to-voice output as an automated measure of intelligibility [18].

However, using a zero-shot ASR model as an evaluator presents a mismatch between the task for which ASR models are trained and the task for which they are being used. Modern ASR models are trained to accurately transcribe speech from large-scale healthy speech datasets, and their performance depends on the similarity between the input speech and the distributions represented in their training data [19]. Brain-to-voice output, in contrast, can contain atypical acoustic characteristics that differ substantially from typical speech [6]. Under such conditions, ASR models may produce errors that differ from those made by human listeners. In particular, Whisper has been shown to generate hallucinated phrases that have no basis in the input audio, with hallucinations occurring under conditions including non-speech audio and low-quality input [20,21]. The accuracy-maximizing objective of traditional ASR also presents the risk that these models could outperform human listeners [22], and thus not reflect the performance of a voice neuroprosthesis in real-world use, where what matters is how well other humans (not machines) can understand the user. This raises an important question for brain-to-voice evaluation: although zero-shot ASR can provide an automated transcription, does its WER reflect how intelligible the speech is to a human listener?

Despite these limitations, pretrained ASR models provide a strong backbone for automated synthesized speech transcription and can be adapted to speech distributions that differ from those represented in their original training data. Pretrained ASR models have been fine-tuned to improve recognition across a range of specialized speech domains, including child speech [23,24], low-resource languages [25,26], and other atypical speech conditions [22,27]. Parameter-efficient fine-tuning (PEFT) methods provide a practical approach for adapting large pretrained models while updating only a small fraction of their parameters, reducing memory, storage and training data requirements compared to full fine-tuning [28–31]. Low-rank adaptation (LoRA) is a widely used PEFT approach that introduces trainable low-rank matrices into selected model layers while keeping the original model weights frozen [32]. For example, LoRA has been successfully applied to Whisper for adapting ASR to atypical speech [33,34] and low-resource languages [26], demonstrating that specialized speech recognition behavior can be learned with substantially fewer trainable parameters. These properties make LoRA well-suited for adapting a pretrained ASR model to the acoustic characteristics of brain-to-voice speech using a relatively small amount of human transcription data.

Here, we develop a human-calibrated ASR pipeline for automated intelligibility evaluation of brain-to-voice BCI output. We collected human transcriptions for ∼14.5 hours of intracortical decoder-synthesized speech (∼72 hours of transcribed speech), which to our knowledge is the largest such dataset to date. We fine-tuned Whisper-large-v3 using LoRA on brain-to-voice speech paired with human transcriptions, with the goal of calibrating model WER to match human transcription WER (Fig. 1). We evaluated the resulting model across speech synthesized by different brain-to-voice decoders and from different acoustic representations, including conditions not encountered during fine-tuning. Fine-tuning increased the correlation between model WER and human WER, and brought the model’s insertion, deletion and substitution error rates closer to those of human listeners. Finally, we show that close agreement with human WER can be achieved using approximately 1.5 hours of brain-to-voice speech for fine-tuning. Together, these results demonstrate a scalable approach for automated intelligibility evaluation that reflects human transcription more closely than zero-shot ASR does.

**Figure 1:**
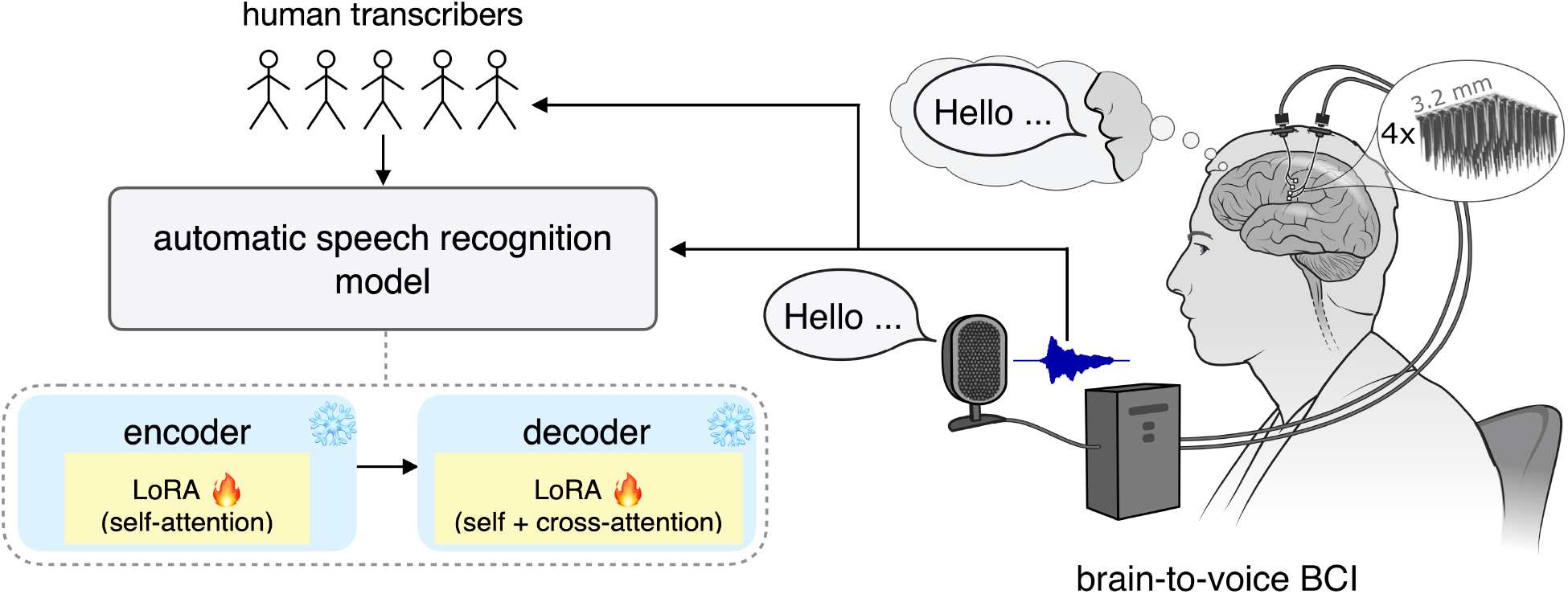
Human-calibrated speech recognition model for speech BCI. A brain-to-voice brain–computer interface (BCI) continuously decoded voice from intracortical neural signals recorded from the speech motor cortex as a participant attempted to speak. This synthesized speech was transcribed by human listeners, providing reference transcriptions that capture how humans perceive the speech. These paired synthesized speech and human transcriptions were used to fine-tune an automatic speech recognition model (Whisper-large-v3) to the characteristics of brain-to-voice speech and human perception. The model’s pretrained weights are frozen, while low-rank adaptation (LoRA) modules added to the attention layers are trained.

## Methods

### Brain-to-voice data corpus

The synthesized speech used in this study was generated using previously developed brain-to-voice decoders: brain-to-voice 1.0 (‘b2v1’) [6] and brain-to-voice 2.0 (‘b2v2’) [7]; no neural data were analyzed directly. Both decoders were developed using intracortical neural activity recorded from participant T15 in the BrainGate2 clinical trial (ClinicalTrials.gov: NCT00912041), which were publicly released in de-identified form [6]. T15 has four Utah arrays implanted in the speech motor cortex [3]. Briefly, both decoders used transformer-based neural networks to map intracortical neural activity, recorded while the participant attempted to speak prompted sentences, to continuous acoustic features that were subsequently converted to voice using a pretrained LPCNet vocoder [35]. The b2v2 decoder additionally used a multitask architecture with multiple output heads and loss functions, including a continuous-output head and a head that predicts discrete acoustic tokens. We refer to speech synthesized from the discrete acoustic tokens as the “alternative acoustic features” condition (Fig. 2). Only speech synthesized from continuous acoustic features was used for fine-tuning the ASR model; speech synthesized from discrete acoustic tokens was used only for evaluation.

**Figure 2:**
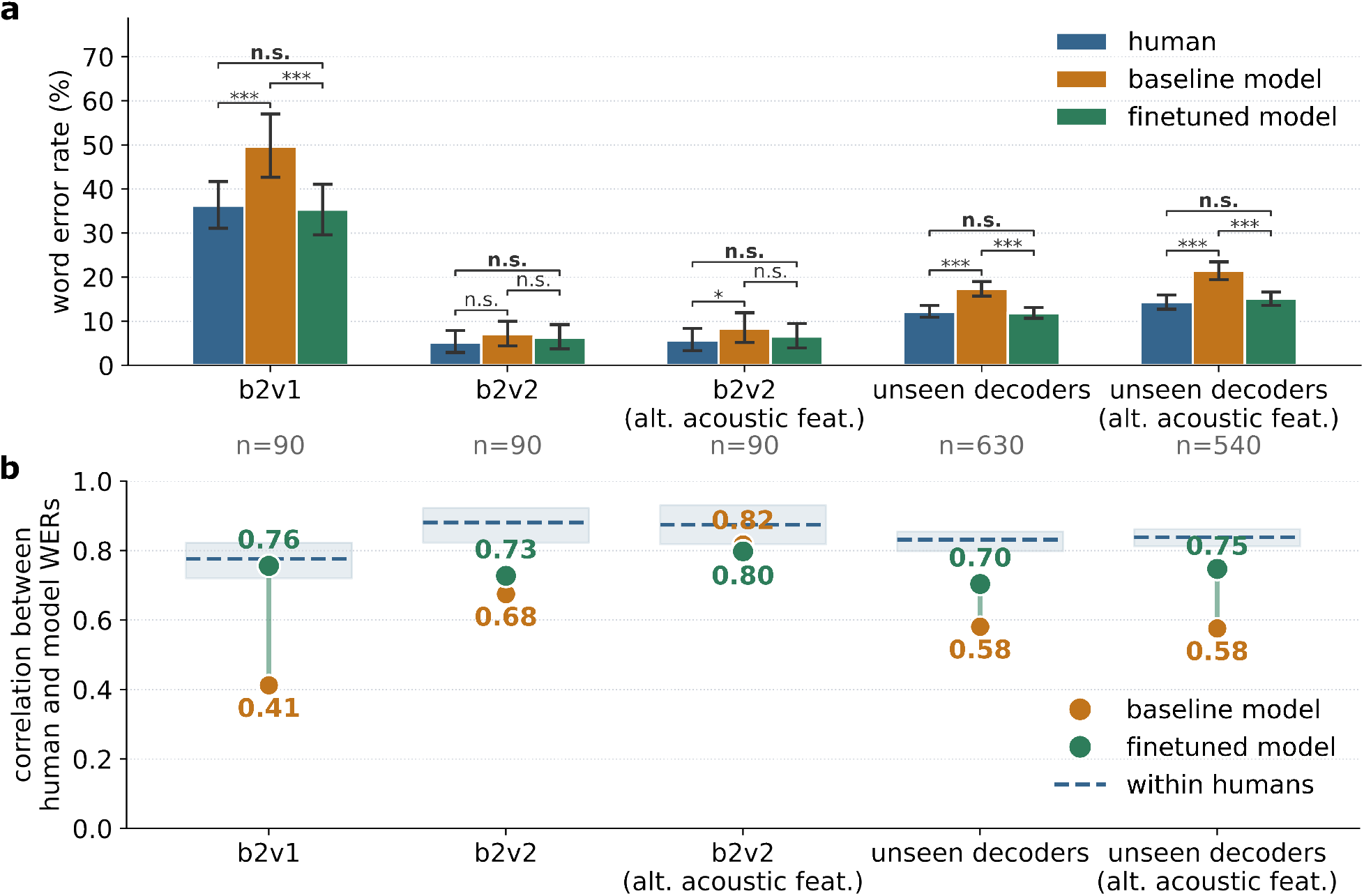
Fine-tuned model yields WERs more correlated with human transcribers. a) WERs for various brain-to-voice evaluation sets. The bars show the mean WER with error bars indicating 95% bootstrap confidence intervals (CI). There is no significant difference between fine-tuned model WERs and human WERs, but there is a significant difference between baseline model WERs and human WERs for most sets (two-sided Wilcoxon signed-rank test with Bonferroni correction; *** *p* < 0.001, * *p* < 0.05, n.s. non-significant). b) Correlation coefficient between model and human WER before and after fine-tuning; correlation increases for almost all decoder versions after fine-tuning. The within-human correlation ceiling (split-half WER correlations with Spearman–Brown correction) is shown as dashed lines with 95% CI as shaded region.

For the primary task of fine-tuning an ASR model, we used the trained b2v1 and b2v2 decoders to synthesize speech from attempted-speech trials in the predefined training split. The fine-tuning dataset comprised 3.6 hours (752 utterances) of synthesized speech from b2v1 and 3.5 hours (698 utterances) from b2v2. To evaluate the ASR models, we synthesized speech using the same decoders on the predefined evaluation split of 90 held-out trials (∼23 minutes). In addition to these two decoders, we also evaluated ablated versions of the b2v2 decoder [7], referred to as “unseen decoders” in Fig. 2. These ablated b2v2 decoders had incrementally increasing numbers of components in the multitask architecture. Speech synthesized by these ablated decoders was not used for fine-tuning the ASR model. In total, we evaluated five groups of brain-to-voice outputs: (1) b2v1; (2) b2v2 synthesized from continuous acoustic features; (3) b2v2 synthesized from discrete acoustic tokens; (4) synthesized speech from all seven b2v2 decoder ablations using continuous acoustic features; and (5) synthesized speech from six ablations using discrete acoustic tokens. The pooled evaluation set therefore comprised 16 decoder–acoustic feature conditions amounting to 1440 utterances (∼6.2 hours).

### Human intelligibility

Each synthesized utterance was independently transcribed by five unique human transcribers using GoTranscript (gotranscript.com), an online transcription service whose transcribers undergo language-specific testing and an application review process before approval. Transcribers were instructed to transcribe the words they perceived in the synthesized speech. When a word was unintelligible, transcribers were instructed to enter “?”. Transcriptions were not required to be grammatically correct. The transcribers were blinded to the prompted sentence and could replay the synthesized speech any number of times when transcribing. No identifying information about the transcribers was collected.

### ASR model selection

We evaluated several pretrained ASR models using zero-shot transcription to identify an appropriate backbone for human calibration (Supp. Fig. 2). Specifically, we tested Whisper-small, Whisper-medium, and Whisper-large-v3 [19] (Throughout this manuscript, ‘Whisper’ refers to Whisper-large-v3 unless otherwise specified), as well as the NVIDIA Parakeet (parakeet-ctc-0.6b) [36], Canary (canary-1b) [37], and Canary-Qwen (canary-qwen-2.5b) models. Whisper-large-v3 was selected as the backbone because it showed the lowest zero-shot WER and the highest correlation with human WER among the models evaluated (Supp. Fig. 2).

### Parameter-efficient fine-tuning of ASR model

We used Whisper-large-v3 as the pretrained ASR backbone. The pretrained model was fine-tuned using low-rank adaptation [32], which keeps the pretrained model weights frozen while learning low-rank updates to selected model weight matrices. LoRA adapters were applied to the query, key, value and output projections of the encoder self-attention, decoder self-attention and decoder cross-attention layers, with a rank of 16 and a scaling factor (α) of 32 (Fig. 1). This LoRA configuration involved training ∼15.7M parameters (∼1% of the 1.55B-parameter Whisper model).

Because human transcribers marked unintelligible words with “?”, we introduced a dedicated <*unk_word*> token into the Whisper tokenizer. During training, standalone “?” markers in human transcriptions were replaced with this token. The tokenizer vocabulary was expanded accordingly, and Whisper’s input embedding and language-model output matrices were resized to accommodate the new token. Under LoRA fine-tuning, only the embedding and output rows corresponding to the newly added <*unk_word*> token were additionally made trainable; gradients for all other vocabulary rows were masked.

The model was fine-tuned using the synthesized utterances from the training splits of the b2v1 and b2v2 decoders. The utterances were divided into training (85%) and validation (15%) splits, stratified by human WER using a fixed random seed. During fine-tuning, five utterance– transcription pairs were created for each synthesized utterance by treating each of its five human transcriptions as a separate training example, resulting in five training pairs per utterance. The model was trained for up to 2,000 steps, with early stopping based on validation-split WER, using the AdamW optimizer with an initial learning rate of 3 × 10^-5^, a linear learning-rate schedule, and a batch size of 32. The checkpoint with the lowest WER against the human reference transcriptions on the validation split was retained. The model was evaluated using greedy decoding. Fine-tuning was implemented in Python 3.10 using the Hugging Face Transformers and PEFT libraries and performed on a single NVIDIA RTX 5090 GPU.

### Word error rates and statistical analysis

WER was calculated with respect to the prompted sentence that the participant attempted to speak. In human transcriptions, each standalone “?” marker was treated as a substitution error for the corresponding word in the prompted sentence. For each neurally synthesized utterance, we first calculated the WERs of the five independent human transcriptions. The median of these five WERs was used as the human WER for that utterance. Model WER was calculated separately for the same utterance, allowing direct utterance-level comparisons between model and human performance. Dataset-level WER was measured as the mean of the per-utterance WERs.

To estimate the within-human WER correlation, we randomly divided the five human transcriptions available for each utterance into two groups. For each utterance, we calculated the median WER within each group and then computed the correlation between the resulting utterance-level WER values across the two groups. We repeated this procedure 100 times using different random partitions and reported the mean split-half correlations with Spearman–Brown correction.

For aggregate WER comparisons (Fig. 2–3), 95% confidence intervals were measured using 10,000 bootstrap iterations with replacement. Pearson correlation coefficients were measured between utterance-level WERs for each pair of evaluation groups: human, baseline model, and fine-tuned model. For comparisons between paired groups, statistical significance was assessed using the two-sided Wilcoxon signed-rank test. *p*-values were corrected for multiple comparisons using the Bonferroni method. Insertion, deletion, and substitution error rates were calculated using the human transcription with the median WER for each utterance.

To further quantify the model and human transcription errors, we identified error positions separately for each synthesized utterance. For each word position, we classified the transcribed word as human error if more than 50% of human transcribers transcribed it incorrectly. We then identified the word positions at which the ASR model made errors relative to the prompted sentence. For each utterance, we measured the Jaccard index between the set of human error positions and the set of model error positions. We compared Jaccard indices between the baseline and fine-tuned models across utterances using a two-sided Wilcoxon signed-rank test.

### Dataset-size analysis

To determine the amount of human-transcribed, neurally synthesized speech required to calibrate the ASR model, we fine-tuned separate models using subsets of utterances from the training split, corresponding to 0.12–4 hours of synthesized speech (Fig. 4). For each target duration, we trained ten models using independently sampled subsets of training utterances. Each subset retained all five human transcriptions associated with every selected utterance. All models were trained and evaluated as described above. We then measured the Pearson correlation coefficient between model WER and human WER on the pooled evaluation set. For each target duration, we report the mean ± standard deviation correlation across ten models.

### Electrode-drop analysis

To evaluate model performance across a broader range of brain-to-voice intelligibility levels, we generated speech using b2v2 decoders trained on data from progressively fewer intracortical electrodes [7] (Fig. 5). For each electrode-count condition, we randomly sampled a subset of the 256 electrodes used by the original b2v2 decoder. Electrode counts ranged from 16 to 256 in increments of 16. A new b2v2 decoder was trained for each electrode subset, and speech was synthesized from the same 90 held-out evaluation trials. The resulting synthesized speech was evaluated using the fine-tuned Whisper model. Human transcriptions were additionally obtained for the 64-, 128-, 192-, and 256-electrode conditions, allowing direct comparisons between model and human WER across intelligibility levels.

## Results

We first characterized variability in human transcription by comparing WERs across transcribers of the same synthesized utterance and examining pairwise correlations between transcribers who shared at least 10 utterances (Supp. Fig. 1). WERs varied substantially among transcribers across all intelligibility levels (Supp. Fig. 1a). Even for synthesized utterances with a median human WER of 0%, individual transcriber WERs ranged from 0–100%. Across transcriber pairs, the median Pearson correlation between their WERs was 0.54 (interquartile range of 0.29–0.70; 232 transcriber pairs), indicating moderate agreement (Supp. Fig. 1b).

### Fine-tuned ASR WER tracks human WER across brain-to-voice decoders

We first evaluated six pretrained ASR models: Whisper-small, Whisper-medium, Whisper-large-v3, NVIDIA Parakeet, Canary, and Canary-Qwen, on held-out synthesized utterances without fine-tuning (Supp. Fig. 2). Whisper-large-v3 achieved the lowest zero-shot word error rate of 19.7% and the highest correlation with human WER (r = 0.62) among the models evaluated. We therefore selected it for LoRA fine-tuning.

We next evaluated the fine-tuned model on synthesized speech from various b2v decoders and acoustic features (Fig. 2). The five evaluation sets comprised b2v1; b2v2 synthesized from continuous acoustic features; b2v2 synthesized from alternative acoustic features; and two aggregate groups of unseen b2v2 decoder ablations synthesized from either continuous or alternative acoustic features. Across all individual evaluation sets, the fine-tuned model’s WER more closely followed human WER than did zero-shot WER. For b2v1, where human WER was 36%, zero-shot WER was 50%, whereas fine-tuning reduced model WER to 35%. There was no significant difference between human WER and fine-tuned model WER across all groups (two-sided Wilcoxon signed-rank tests with Bonferroni correction; Fig. 2a).

After fine-tuning, the correlation between model WER and utterance-level human WER increased in four of the five sets, with the largest improvement for b2v1 (0.41 to 0.76; Fig. 2b). For b2v2 syntheses using alternative acoustic features, the correlation remained essentially unchanged (0.82 before versus 0.80 after fine-tuning), as the zero-shot correlation was already near the estimated within-human correlation ceiling. Overall, fine-tuned model correlations approached this ceiling.

### Fine-tuning reduces non-human-like transcription errors

We next examined the relationship between model and human WER for individual synthesized utterances pooled across all evaluation sets (n = 1440; Fig. 3a). Fine-tuning increased the correlation between model and human WER from r = 0.62 to r = 0.76 and reduced pooled model WER from 19.7% to 13.9%, close to human WER (13.7%). Baseline model WERs deviated from human WERs for a subset of utterances, including high-WER outliers associated with repetitive or runaway speech-to-text generations (bottom-right points in Fig. 3a left panel). Following fine-tuning, model WERs shifted closer to the identity line, and these high-WER hallucination outliers were substantially reduced.

**Figure 3:**
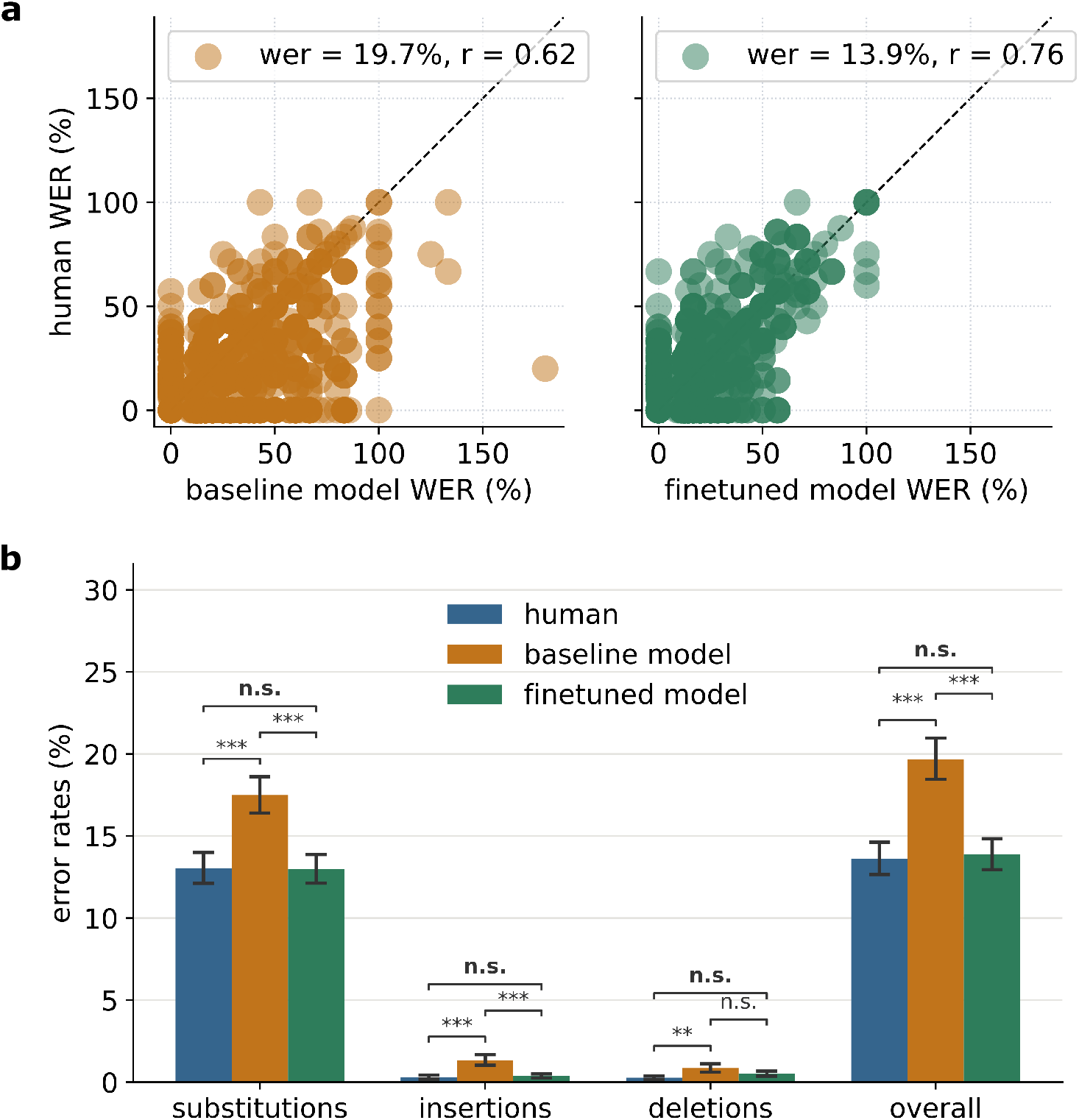
Fine-tuned model error rates and error types are comparable to those of humans. Scatter plot of human WER against baseline and fine-tuned model WER. WERs of all synthesized utterances in the pooled evaluation set are included (n = 1440). The dashed black line is the identity line showing parity between ASR and human WER for an utterance. After fine-tuning, correlation increases and points cluster closer to the parity line, with hallucination-driven outliers (bottom-right of plot) curbed. **b)** Insertion, deletion, and substitution error rates for transcriptions from humans, baseline and fine-tuned model (n = 1440). Bars plot the mean error rates for the evaluation set; error bars denote the 95% bootstrap confidence intervals. Across all error types and overall word error rate, the fine-tuned model had closer WERs to humans (two-sided Wilcoxon signed-rank test with Bonferroni correction; *** *p* < 0.001, ** *p* < 0.01).

We further decomposed WER into insertion, deletion, and substitution errors (Fig. 3b). The baseline model showed higher insertion, deletion and substitution error rates than human transcribers, whereas fine-tuning reduced these error rates toward the corresponding human values. These results indicate that calibration altered not only the overall magnitude of WER but also the relative distribution of insertion, deletion, and substitution errors toward the pattern observed in human transcription. The fine-tuned model’s error positions also overlapped more with human consensus error positions than did those of the baseline model (Jaccard indices of 0.56 and 0.48, respectively; two-sided Wilcoxon signed-rank test, *p* < 0.001). Example transcriptions from human transcribers, baseline model and fine-tuned model are provided in Table 1.

**Table 1:**
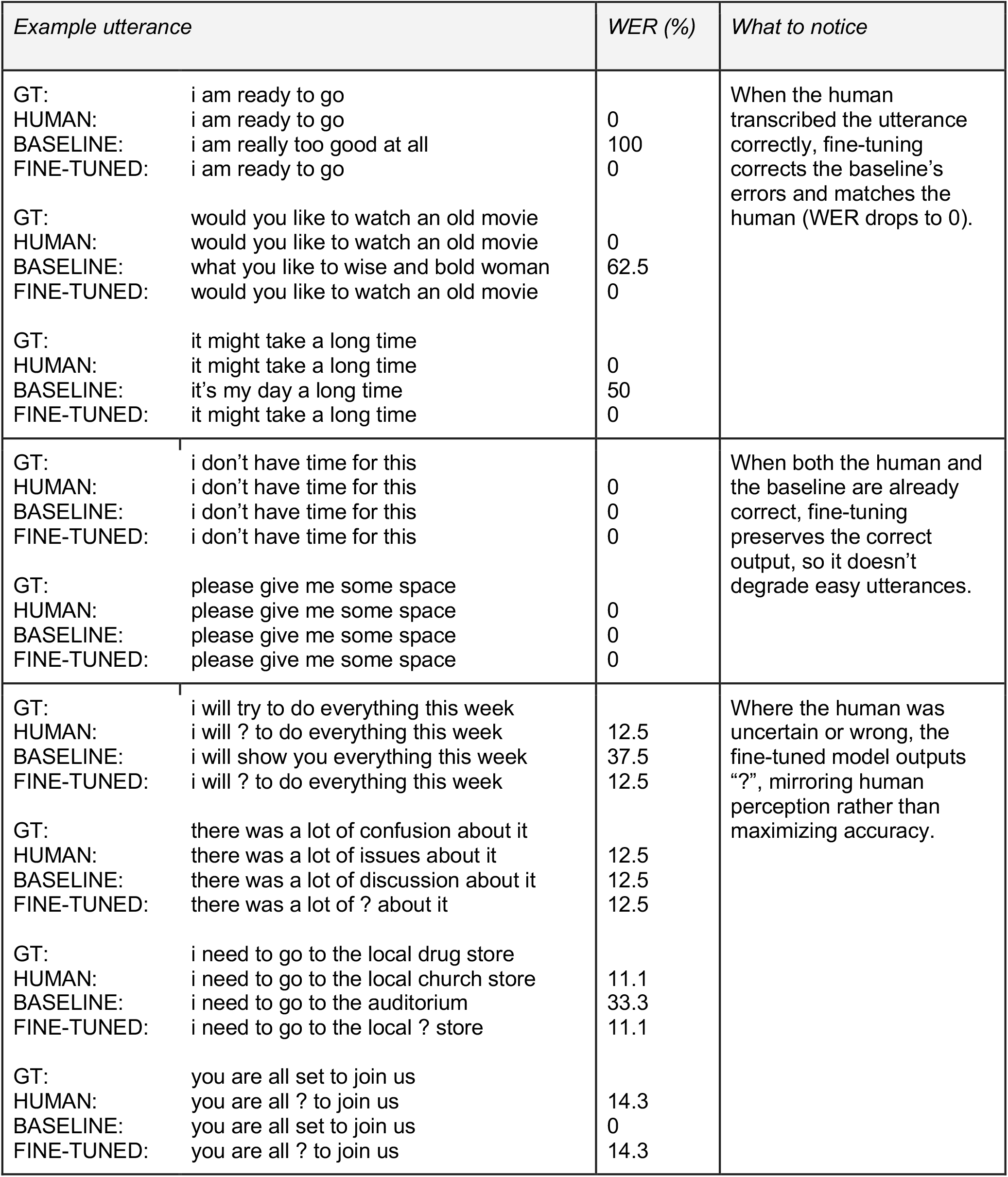
Example transcriptions from human, baseline model and fine-tuned model. GT refers to ground-truth transcript (what the participant attempted to say). HUMAN corresponds to the human transcription with the median word error rate.

### Fine-tuning performance plateaus at approximately 1.5 hours of synthesized speech

We next examined how much human-transcribed brain-to-voice speech data was required to fine-tune the ASR model (Fig. 4). The correlation between model WER and human WER increased with the amount of synthesized speech, reaching r = 0.74 with 1.5 hours of synthesized data. Beyond this point, additional data produced only marginal changes in correlation, suggesting diminishing returns for this model architecture and speech-synthesis distribution. Because each utterance was paired with five independent human transcriptions, the 1.5-hour condition corresponded to approximately 7.5 hours of utterance–transcription training pairs.

**Figure 4:**
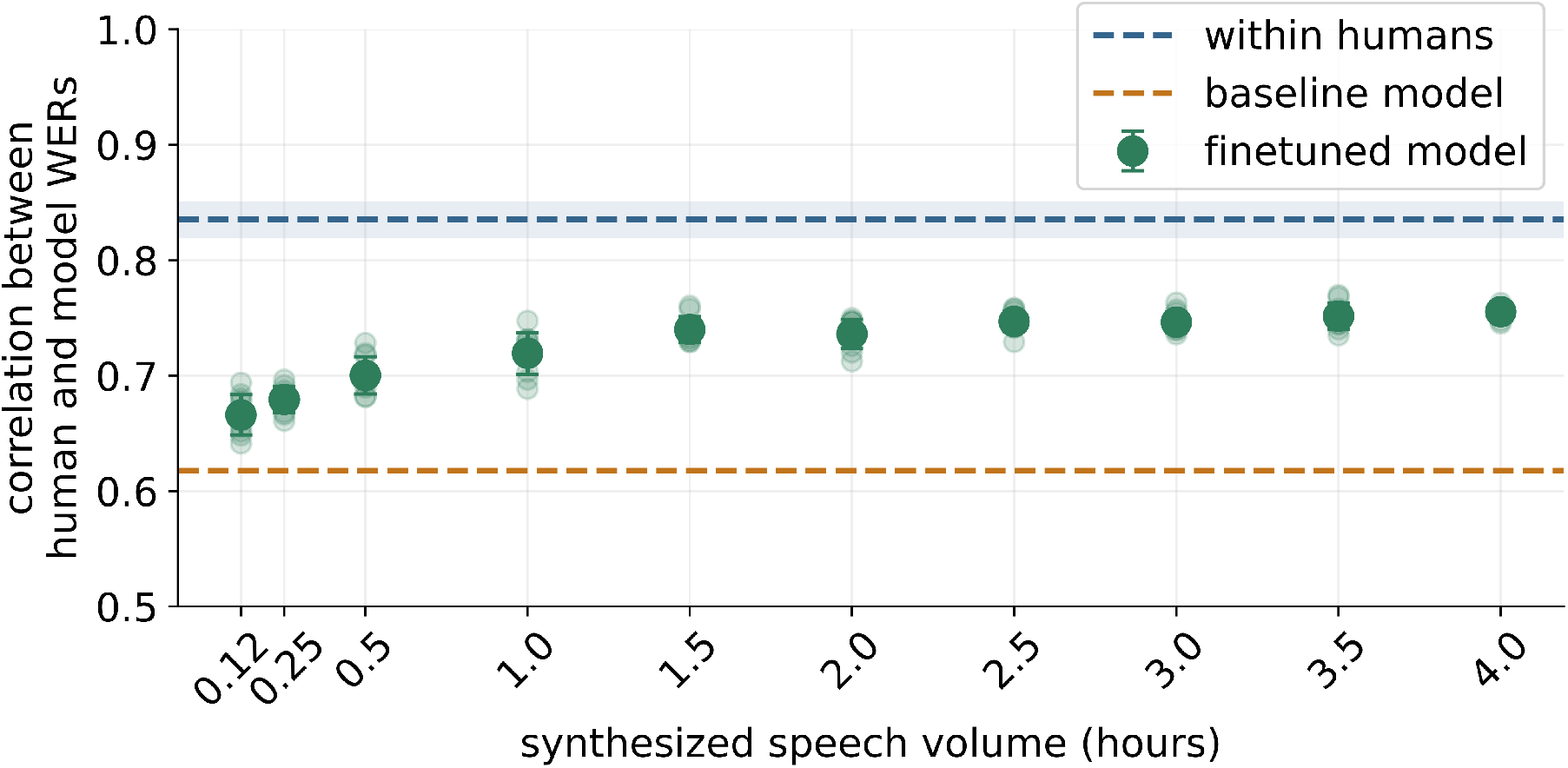
Human-calibrated model performance saturates with 1.5 hours of synthesized speech. The ASR model was fine-tuned using increasing amounts of brain-to-voice synthesized speech to evaluate the amount of data required for effective adaptation. Each model was evaluated on the pooled evaluation set (n = 1440 utterances). The green filled circles with error bars denote the mean ± standard deviation correlation, with individual seed correlation in faint circles.

### Fine-tuned model WER predicts intelligibility across brain-to-voice decoders with various electrode counts

Finally, we demonstrate how a human-calibrated ASR model can facilitate analyses that would be prohibitively time-consuming and labor-intensive to perform using only human transcriptions. Here, we tested whether the fine-tuned ASR model continued to track human WER across a wide range of brain-to-voice intelligibility. We synthesized speech using b2v2 decoders trained with progressively larger subsets of the available intracortical electrodes, ranging from 16 to 256 electrodes (Fig. 5). As expected, fewer electrodes produced a graded range of speech intelligibility.

**Figure 5:**
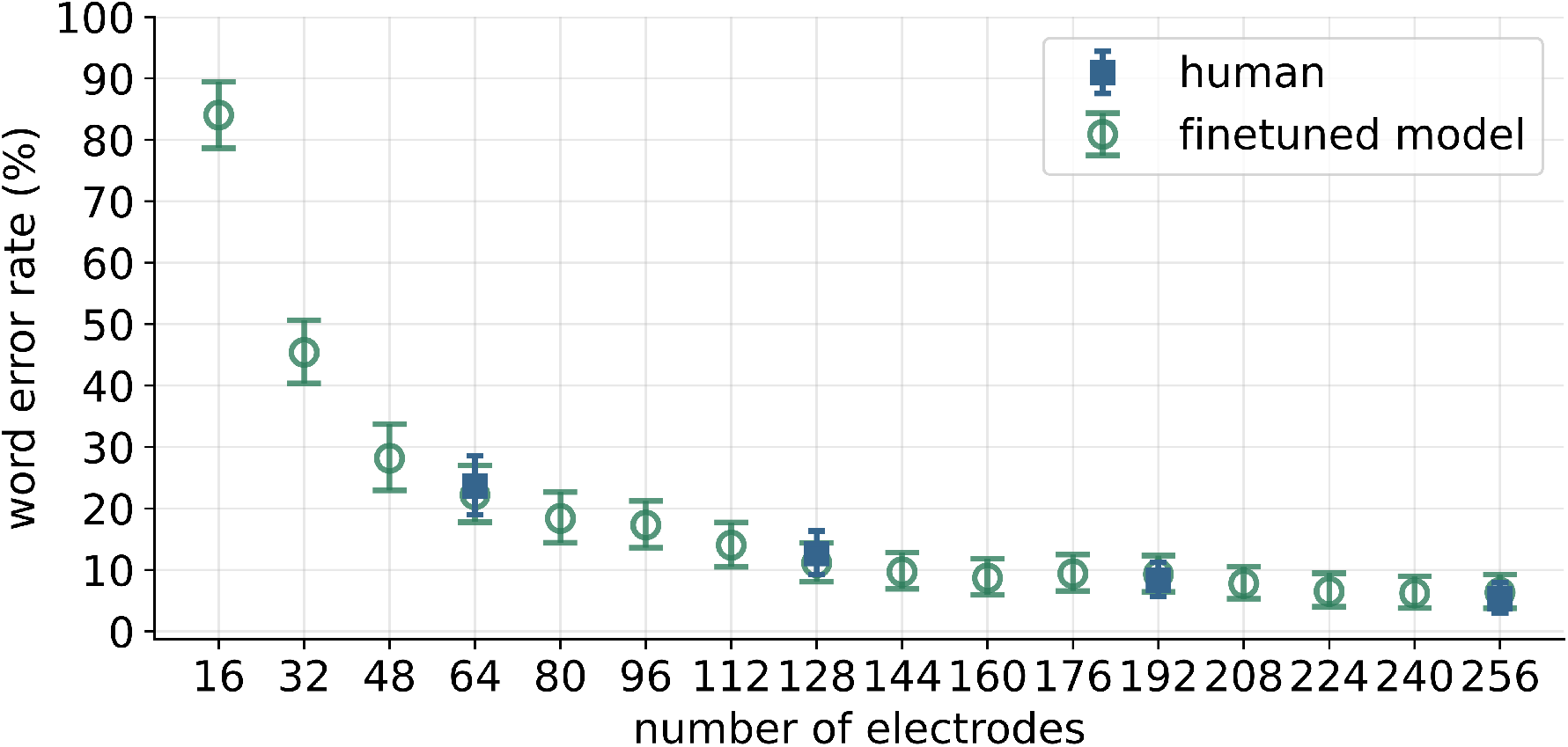
Fine-tuned model WER tracks human WER across decoders with randomly dropped electrodes. Each decoder was trained on a single random subset of 16–256 electrodes (increments of 16) and evaluated on the same 90 held-out trials. Points show mean WER across utterances (n = 90 per condition); error bars are 95% bootstrap confidence intervals. Where human transcriptions were available, the fine-tuned model and human WERs were similar.

Across the electrode-count conditions for which human transcriptions were available, WER measured by the fine-tuned model tracked the changes in intelligibility reflected in human WER. These results indicate that the calibrated model continued to track human WER across a range of experimentally induced reductions in decoder performance. Because obtaining human transcriptions for every decoder configuration is labor-intensive, the fine-tuned ASR model could enable more systematic evaluation of speech intelligibility across larger numbers of decoder configurations.

## Discussion

We developed a human-calibrated automatic speech recognition approach for evaluating the intelligibility of brain-to-voice synthesized speech. Fine-tuning Whisper-large-v3 with LoRA on a modest amount of human-transcribed neurally-synthesized speech produced an automated evaluator whose WER more closely tracked human WER than zero-shot ASR does, including insertion, deletion and substitution error rates. This improvement was observed across brain-to-voice decoder variants.

This work extends prior efforts to use pretrained ASR for automated evaluation of brain-to-voice [18]. Here, we selected an ASR model based on its zero-shot behavior and then adapted it using human-transcribed synthesized speech. More broadly, automated intelligibility assessment of dysarthric or other atypical speech has been developed as classification or regression approaches, predicting a sentence-level intelligibility score on a subjective rating scale. For example, Merler et al. [17] used Whisper as a feature extractor to predict listener effort ratings. In contrast, our approach retains WER as the evaluation metric, quantifying how many of an utterance’s words a listener can correctly identify. This directly links automated brain-to-voice evaluation to the human-transcription-based WER that is standard in assessing synthesized speech [6,38].

The amount of synthesized speech required for fine-tuning was relatively modest. Performance plateaued at approximately 1.5 hours of unique synthesized speech. This is encouraging given that intracortical brain-to-voice datasets are typically limited [6]. The electrode-dropping experiments further showed that the fine-tuned model continued to track human WER across a range of intelligibility levels induced by reducing the number of electrodes. Together, these results suggest that human-calibrated ASR could reduce the amount of manual transcription required when evaluating many decoder configurations.

Our findings should nevertheless be interpreted within several limitations. First, all data used for calibration and evaluation were generated by intracortical brain-to-voice decoders developed for a single BrainGate2 clinical trial participant. We focused on this dataset because it demonstrated high-quality speech synthesis and because, to our knowledge, comparable brain-to-voice BCI outputs from other systems were not publicly available. Whether calibration learned from one participant and decoder family generalizes to synthesized speech from other systems remains unknown. Importantly, generalization also depends on the human transcription reference: different listener populations might transcribe the same synthesized speech differently. Continued adaptation and validation using human-transcribed speech from diverse neural decoders and listener populations will be important for establishing broader generalizability. Second, decoder performance can vary over time as neural signals and decoding models change across recording sessions. Our experiments do not capture this longitudinal variation, and longitudinal brain-to-voice datasets spanning many sessions will be needed to determine whether calibration remains aligned with human transcription over time. To our knowledge, such datasets are not yet available. Finally, human transcription provides an operational rather than absolute reference for intelligibility, as listeners may differ in how they transcribe ambiguous or unintelligible speech. Moreover, this initial ASR evaluation tool, like our human transcription dataset, focuses only on WER as a metric of intelligibility, which we view as the baseline requirement for a voice neuroprosthesis. However, there are other important aspects of voice quality that future work should develop methods for automatically quantifying.

To facilitate broader neural-speech synthesis model evaluation and development, we will make the human-transcribed synthesized speech dataset used in this study (∼14.5 hours of synthesized speech; ∼72 hours of transcribed utterances) publicly available upon peer-reviewed publication. These data will provide a starting point for improving automated evaluators calibrated to human transcription. We encourage the voice neuroprosthesis community to build on these resources and contribute additional datasets of human-transcribed synthesized speech (from other participants, recording modalities, decoder architectures etc.). Aggregating such data will enable continued adaptation of the ASR model toward a more generalizable automated evaluator that can be applied across increasingly diverse speech neuroprostheses.

An important future direction is to determine whether the calibrated ASR can be incorporated into brain-to-voice decoder development. For example, it could potentially provide a monitoring measure or serve as a loss function to optimize brain-to-voice decoders, or support evaluation of large numbers of decoder configurations (e.g., hyperparameter searches). Such applications would need prospective validation, particularly to ensure that optimizing against the automated evaluator does not introduce systematic differences between model-based evaluation and human perception.

## Conclusion

We developed a human-calibrated automatic speech recognition approach for scalable and reproducible evaluation of brain-to-voice speech intelligibility. By fine-tuning a pretrained ASR model with a modest amount of human-transcribed brain-to-voice speech, we produced an automated evaluator whose WER more closely tracked human-derived WER across brain-to-voice decoder variants and levels of intelligibility than zero-shot ASR does. Rather than optimizing for conventional transcription accuracy, this approach adapts the evaluator to the characteristics of neurally-synthesized speech while retaining WER as a familiar and interpretable metric. By reducing reliance on manual transcription, this pipeline provides a scalable complement to human evaluation and could facilitate more rigorous and systematic evaluation of voice neuroprostheses.

## Data and code availability

Data and code will be shared publicly upon acceptance of the manuscript.

## Acknowledgements

This work was supported by the UC Davis School of Medicine Cultivating Team Science Award, NIH/NCATS: UC Davis Clinical and Translational Science Center (CTSC UL1-TR001860), and a Google Research Scholars award to S.D.S. We thank Daniel C. Comstock for providing feedback on the overall study design and assisting with the pilot evaluation of brain-to-voice 1.0.

## Supplementary information

**Supplementary Figure 1:**
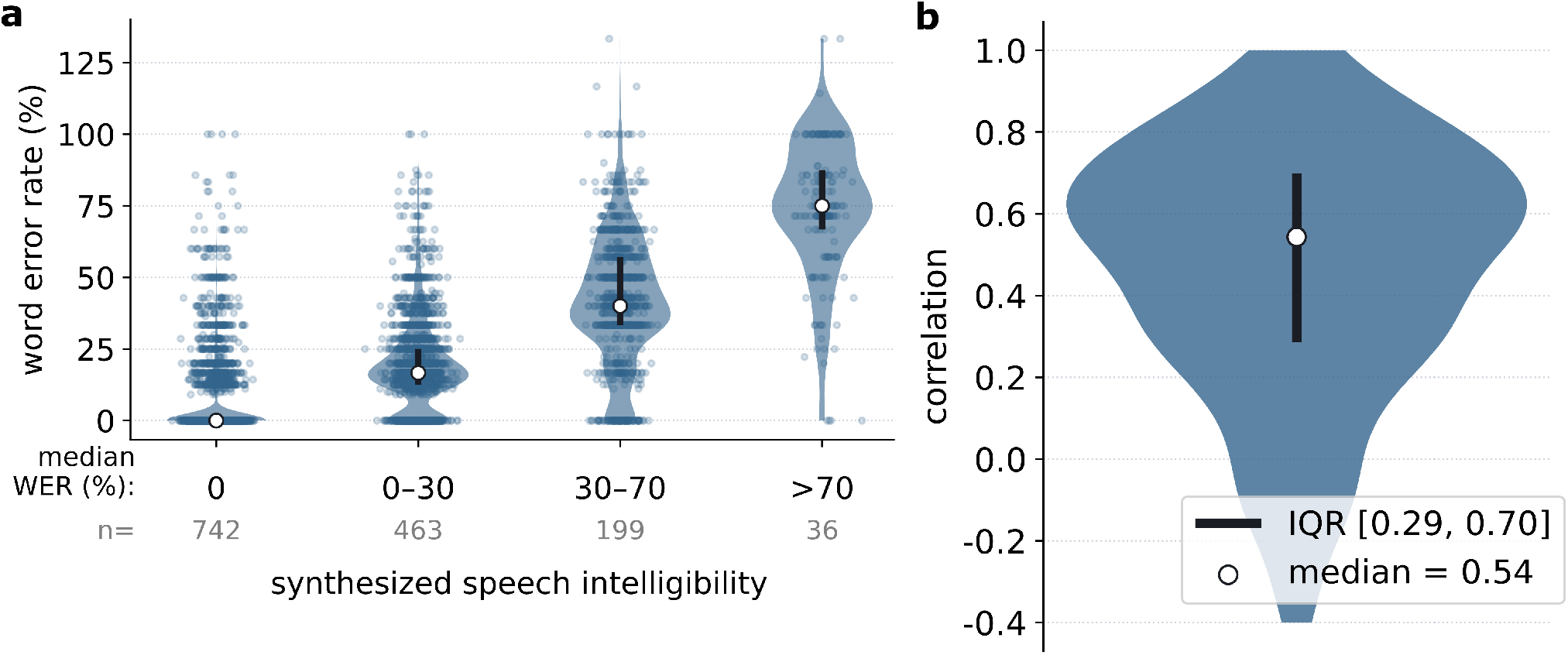
Human transcription overview. a) WERs of individual transcribers for synthesized utterances, grouped by intelligibility. WERs varied widely across transcribers in all groups. b) Distribution of pairwise Pearson correlation between WERs of transcriber pairs who transcribed at least 10 shared utterances. There is moderate agreement between transcribers but with high variance in transcription accuracy.

**Supplementary Figure 2:**
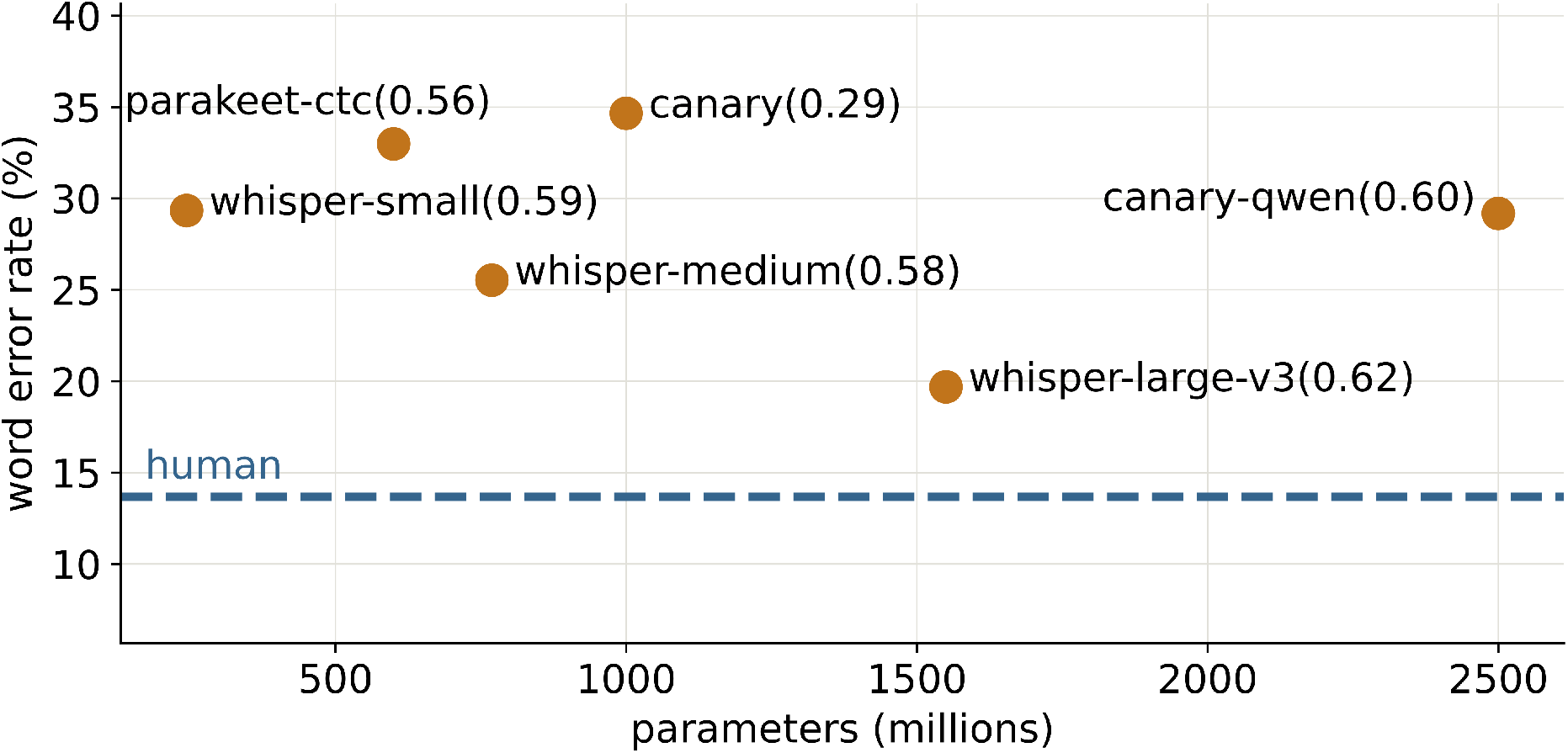
Zero-shot word error rates of open-source automatic speech recognition models. Each point shows a pretrained model’s mean WER on the pooled evaluation set (n = 1440 utterances), plotted against its total number of parameters. Values in parentheses are Pearson correlation coefficients between human WER and model WER across utterances.

